# Stand age drives changes in the biodiversity and microclimate of oil palm plantations

**DOI:** 10.1101/2025.10.31.682997

**Authors:** T.A. De Lorm, M. Barclay, H. Bernard, E.R Bush, C. Carbone, S.R. Hardwick, S. Milne, J.M. Parrett, M. Pfeifer, J.M. Rowcliffe, A. Sharp, E. Slade, O. Wearn, R. M. Ewers

## Abstract

Tree plantation age strongly dictates its environment. As they mature, the canopy closes and habitat complexity tends to increase, which in turn affects biodiversity. However, for oil palm, one of the most important and widespread tree plantations, we lack a detailed understanding of how this process affects microclimate and biodiversity. We therefore compiled environmental and biodiversity data of oil palm plantations that were repeatedly sampled over a 6 year period in Malaysian Borneo. Using these data, we made a chronosequence of the microclimate and biodiversity of oil palm plantations between the ages 5 and 18. We found that oil palm plantations increasingly buffer temperature and humidity over time, becoming cooler and less dry. Animal responses differed between taxa: spider and beetle abundance increased, ant abundance decreased, and mammal and dung-beetle abundances stayed constant. The diversity of beetles in general increased and family-level community composition changed, while dung-beetle and mammal diversity stayed constant. Some taxa are therefore likely limited by the harsh microclimate of young plantations’, while increased habitat complexity and fruit and leaf production in older plantations could also drive increased abundances of other taxa. Our results reveal that oil palm plantations undergo considerable changes, even over a decade after establishment. Correspondingly, plantation age should be considered in studies into oil palm plantations’ impact on climate and biodiversity, and landscape level diversity could be increased by allowing multiple oil palm stand-ages to co-exist.

## Introduction

Throughout the life cycle of a tree plantation its environment and biodiversity undergo substantial changes, in some ways resembling the ecological succession of natural forests (Pashkevich et al., 2021). The establishment of tree plantations has had a big impact on biodiversity, as they cover more than 230 million hectares (Richter et al., 2024), and their biodiversity tends to be lower than the natural habitats they often replace (Fitzherbert et al., 2008). However, plantation structure, microclimate, and microclimate availability is influenced by stand-age (Luskin & Potts, 2011; Santoandré et al., 2019). Understanding how tree plantations’ biodiversity develops through time is therefore important to evaluate their overall impact on biodiversity.

Oil palm (*Elaeis guineensis*) is one of the most widespread and rapidly expanding tree crops in the tropics (Phalan et al., 2013). In the two largest national producers, Indonesia and Malaysia, respectively 8% and 18% of the countries’ land area was used to grow oil palm in 2020 (Badan Pusak Statistik, 2021; Ghulam Kadir, 2022), and it is also gaining footholds in Africa and South America (Ordway, Asner & Lambin, 2017; Furumo & Aide, 2017). One of the reasons for its ubiquity is its high land-use efficiency (Carter et al., 2007), with yields averaging 2.91 tonnes per hectare, approximately four times more than sunflower, the second highest yielding oil crop (Ritchier & Roser, 2021). As a result, oil palm delivers 36% of the global vegetable oil on less than 9% oil crop producing land (Ritchier & Roser, 2021).

Due to its rapid expansion, the palm oil industry has been subject to substantial public scrutiny, as it has been linked to deforestation and biodiversity loss (Pendrill et al., 2019). Indeed, palm oil plantations are drastically less biodiverse than tropical forests ((Meijaard et al., 2020)). The abundance and diversity of nearly all arthropod taxa are reduced, and their community composition changes (Turner & Foster, 2009; Savilaakso et al., 2014). The diversity of beetles in general (Chung et al., 2000) and dung beetles in particular (Davis & Philips, 2005) are lower compared to primary forests, and canopy spider adundance, biomass and species richness are twice as low in oil palm plantations (Ramos et al., 2022). Mammals are similarly severely affected (Daniel et al., 2022). Wearn et al. (2017) found a 47% decrease in mammal species abundance in oil palm plantations compared to forest habitat, and most species that remain are unlikely to form self-sustaining populations in plantations. Some native mammals do occur in plantations, such as bearded pigs *Sus barbatus*, and mesopredators like the leopard cat *Prionailurus bengalensis,* which appear to benefit from the high abundance of rats (Harich & Treydte, 2016; Davison et al., 2019).

The biodiversity of tree plantations, however, is not constant througout their lifecycle, as the environmental conditions of tree plantations vary considerably through time (Pashkevich et al., 2021). Newly established plantations are open habitats, but as plantations age, canopy height and cover increases, causing microclimatic conditions to increasingly resemble those of forests, by buffering temperature and humidity (Luskin & Potts, 2011; Santoandré et al., 2019, Hardwick et al., 2015). Correspondingly, in Uruguan and Brazilian pine plantations, Santoandré et al. (2019) showed that ant assemblages became more similar to natural forests as plantations aged, and less similar to grassland communities. Flying beetle diversity is higher in mature pine plantations than young plantations in New Zealand (Hutcheson & Jones, 1999). Mammal species richness in Urugayan eucalyptus plantations peaked at intermediate plantation age (Cravino, Andrés Martínez-Lanfranco & Brazeiro, 2023), while plant diversity steadily declined along a chronosequence of Argentinian eucalyptus plantations (Pairo et al., 2021). Songbird diversity similarly peaked at intermediate ages of seral tree plantations in Oregon, USA, but decreased after canopy closure (Harris & Betts, 2021). These studies reveal that biotic communities change across a plantation’s life cycle, but that biodiversity does not necessarily increase, since responses vary amongst crop-types and taxa.

Most studies focussed on oil palm’s microclimate and biodiversity have treated plantations as a static environment. Two notable exceptions are Luskin & Potts (2011) and Pashkevich et al. (2021). Luskin & Potts (2011) measured the microclimate of 8 and 22 year-old plantations in Peninsular Malaysia. They found that the microclimate became more strongly buffered under old palms, although even old plantations remained significantly hotter and drier than intact forests during the day. Overall, older plantations were 2.84°C hotter than forests, and the young plantations were 1.20°C hotter than old plantations. Nighttime plantation temperatures, however, did not differ from forests. Pashkevich et al. (2021) assessed the arthropod abundance and order-level diversity of 31-to 33-year-old first-generation oil palm plantations, and 1, 3, and 8 years after replanting. The arthropod communities of 31-to 33 year-old and 8-year-old second-generation palms were most alike. The community composition of 1- and 3-year-old replanted palms significantly different from these, which was mainly driven by lower ground Coleoptera, Dermaptera, and Lepidoptera abundances. However, their abundances bounced back to the levels of the mature plantations within 3 years after replanting. These studies indicate that the environment and arthropod communities of oil palm plantations undergo changes, both after the establishment of first-generation plantations, and after replanting.

Still, we lack a thorough understanding of how the environment and biodiversity of oil palm plantations develop as plantations mature. Although Luskin & Potts (2011) showed that plantations become cooler over time, they only looked at 8- and 22-year-old plantations, so we lack an understanding of how the microclimate develops between and beyond those ages, as plantations are typically replanted after 25 to 30 years. The age at which a plantation’s microclimate stabilises therefore remain unknown. Furthermore, to the best of our knowledge, no studies have looked at how the biodiversity of first generation oil palms develops over time.

Understanding how the biodiversity in an oil palm plantation varies through its lifecycle can help guide landscape planning. If plantations exhibit large temporal variation in their environmental conditions and biotic communities, ensuring that landscapes contain multiple stand ages can increase landscape-scale habitat heterogeneity (Luskin & Potts, 2011). Luskin & Potts (2011) further highlight that if certain species only occur in specific stages of the lifecycle of a plantation, ensuring that strips of this age-class exist between natural habitats can increase habitat connectivity.

Here, we study a chronosequence of the microclimate, and abundance of mammals, beetles, ants and spiders, and the diversity of mammals and beetles in oil palm plantations. We used data collected at the SAFE Project in Malaysian Borneo (Ewers et al., 2011), a site that includes oil palm plantations of different ages that have been studied for over a decade, allowing us to compile data from 5- to 17-year-old plantations. In this way, we sought to improve our understanding of how the environment and biodiversity of oil palm plantations change over time, by quantifying the changes in temperature, humidity, and invertebrate and mammal diversity and abundance.

## Methods

### Field site

All data were collected at the Stability of Altered Forest Ecosystems (SAFE) Project landscape in Sabah, Borneo (4.43° N, 117.35° E, Ewers et al., 2011). We used data from oil palm plantations and a primary forest in the Brantian-Tantulit Virgin Jungle Reserve, the edges of which have undergone light illegal logging (figure S1). The SAFE Project included three plots across two oil palm plantations, one of which was established in 2000 (4°63’80.1, 117°45’36.7), and two of which were established in 2006 (4°65’45.8 N, 117°45’36.0° E and 4°64’62.8, 117°43’99.1). The elevation of sample sites ranged from 256 to 599 m. Sampling followed a fractal spatial design. Each fractal order consisted of triangles of sampling points, whose centres formed the vertices of the next fractal order (figure S1). This pattern is repeated for a total of 4 fractal orders (for a detailed description of the sampling design, see Ewers et al., 2011). The closest forest to these plantations is a logged forest, from which no data were collected, as it consists of difficult to reach terrain. The distance between the oil palm sites and this logged forest ranged from 0.23 to 2.32 km. The distance to the primary forest at which data was collected ranged from 7.2 to 9.8 km. We collated biodiversity and microclimate data collected at these plantations between 2011 and 2017 (Figure S2). This allowed us to make chronosequences of the plantation microclimate and biodiversity, from ages 5 to 17 years.

### Microclimate data

Microclimate data were collected between January 2011 and November 2015 (Hardwick & Ewers, 2018; Hardwick, Nilus & Ewers, 2018; Hardwick et al., 2015). Hygrochron iButtons (Maxim Integrated Systems) were deployed at 59 shaded sites in oil palm plantations, at 1.5 metres height. Temperature and relative humidity (temperature accuracy <±0.5 °C, RH accuracy <±5%) readings were taken every 3 hours starting at noon, resulting in eight datapoints per day. To compare the plantations microclimate to an intact forest, we used data collected at 18 sites in the nearby old growth forest, using the same methods.

### Invertebrate traps

Between 2011 and 2017, a total of 275 combined malaise / flight-interception / pitfall-traps were deployed at the SAFE Project’s oil palm sites (Sawang et al., 2019; Sharp et al., 2019). These traps contained a 25 cm diameter top funnel, and a cross-intersecting PVC flight interception vane, to capture flying inverts. To capture ground-dwelling inverts, 20 cm diameter pitfall traps were dug into the ground. For the sampling campaigns between 2011 and 2013, all beetles were identified to family level. In 2012 and 2017, only invertebrates belonging to a small set of major taxonomic groups were recorded (spiders, beetles, and ants). Consequently, our analyses are restricted to these three groups.

### Dung beetle traps

Dung beetle trapping was conducted in February 2011 and February 2015 in the oil palm plantations (Slade et al., 2019a, 2019b; Raine et al., 2018). In both campaigns, 27 pitfall traps filled with a water, salt, and detergent solution, and baited with 25 g of human faeces, were set out for 48 hours. The trap’s contents were preserved in 80% ethanol and frozen, after which all beetles were identified to species level. Specimens were identified using Balthasar (1964), (Boucomont, 1914), the works on Bornean Scarabaeinae by Ochi and Kon (e.g., Ochi & Kikuta, 1996), and the reference collections housed in the Oxford University Museum of Natural History (OUMNH). Voucher collections are deposited at the OUMNH, Universiti Malaysia Sabah, and the Forest Research Centre in Sandakan, Sabah.

### Camera traps

From December 2013 to February 2014, 72 camera traps (Reconyx HC500, Holmen, Wisconsin, USA) were deployed in the oil palm plantations (Wearn et al., 2017). To pick camera trap locations, a 4 by 12 grid with cell size 23 m was overlaid on each side of the second fractal order triangles. On each of the 9 grids, 8 camera traps were then placed on random points of the grid. Photos of mammals were identified and used in the current study. Detections of the same species that were more than 30 minutes apart were treated as independent detections.

### Data analysis

All statistical analyses were conducted using R (version 4.3.0, R Core Team, 2023).

#### Microclimate data

To assess how microclimate is affected by plantation age, we modelled the temperature and humidity of plantations, relative to the nearby old growth forest, as a function of plantation age. For each temperature and relative humidity reading from the plantation sites, we calculated the difference between that reading and the mean of the 18 readings taken at the same timepoint in the old growth forest. Measurements between 09:00 and 18:00 hrs were classified as daytime readings, and measurements between 21:00 and 06:00 hrs were classified as nighttime readings. For each day and night we averaged the difference, so that we ended up with four response variables per 24 hours: the daytime and nighttime average temperature and relative humidity difference. Lastly, we calculated the difference between the maximum and minimum temperature for each 24 hour period.

We fitted separate Generalised Additive Mixed Models (GAMMs) with these five response variables using the MGCV package (Wood, 2023). Each model included plantation age as a smooth response variable, and elevation as a linear variable. For models with temperature as response variable, old-growth forest temperature was included as a linear predictor variable, and for the models with relative humidity as response variable, both the old-growth forest temperature and relative humidity were included as linear predictor variables. We tested whether including the old-growth forest temperature and relatively humidity improved the models using the Akaike Information Criterion (AIC, Akaike, 1973). We used the less complex model if ΔAIC < 3.

To account for spatial autocorrelation, we used sites’ location as a nested random effect, nesting the fractal order sites within each other. We included time as the number of days since the first datapoint as an autoregressive term, to account for temporal autocorrelation. This term included an autoregressive term (p) and the moving average term (q) – i.e. the number of previous observations and previous prediction errors used to predict the current value. The p and q term were fixed at 1, to keep models as simple as possible, while still accounting for temporal autocorrelation and avoiding pseudoreplication.

#### Biotic data

To model how taxon abundances changed with plantation age, we used negative binomial generalised linear mixed effects models using the *glmmTMB* package (Brooks et al., 2023). We used abundance data for all beetles, dung beetles, mammals, ants, and spiders, as number of individuals per trap. For each taxon, we fitted models with plantation age and distance to nearest forest. We fitted various model configurations with age and distance as fixed effects, testing plantation age and distance to forest as linear and quadratic predictors individually, as well as in combined linear and quadratic forms, and as a linear interaction term. We also included a nested random variable with the four fractal orders to correct for spatial autocorrelation. The combined pitfall/malaise/flight-inception traps consisted of separate data for the top and bottom funnels of the traps, so we also included trap-compartment as a fixed effect in these models.

To assess whether community composition changed over time, we used multivariate negative binomial generalized linear models, using the *MVABund* package (Wang et al., 2012). We fitted separate models for the abundance of beetles (family-level), dung-beetles (species-level), and mammals (species-level), and restricted this analysis to these three groups. We fitted models with plantation age and distance to forest as fixed effects. To assess which taxa caused significance of results, we fitted post-hoc univariate ANOVA’s. These bootstrapped ANOVA’s used case resampling with 999 bootstrap iterations, adjusted p-values for multiple testing by step-down resampling, and used plantation number as a block variable.

To test whether species/family diversity changed with plantation age, we also fitted species accumulation curves for (dung-)beetles and mammals, using the *iNEXT* package (Hsieh, Ma & Chao, 2016). For each sample – one trapping occasion for the (dung-)beetles, and one sampling round for the camera trapping data – we rounded off the plantation age to year-level at the time of sampling. For each age, we fitted a separate species accumulation curve. For mammals, we fitted a separate species accumulation curve for each plantation, resulting in three species accumulation curves. For dung-beetle species and beetle families, we fitted species accumulation curves for four plantation ages. All the curves were extrapolated to the same number of samples, which was twice the sampling size of the year/site with the smallest coverage, to allow direct comparisons between years/sites. Estimates were bootstrapped with 999 iterations. To test if species richness significantly differed between years, we looked whether the 84% confidence intervals overlapped, as this has been shown to correspond to using p = 0.05 as cut-off for significance (MacGregor-Fors & Payton, 2013).

## Results

### Microclimate data

For the daytime and nighttime temperature models, including the temperature of the old growth forest improved the model (Table S1). For the daytime and nighttime humidity models, the best fitting models included both the relative humidity and temperature of the old growth forest (Table S1). The best fitting model with daily maximum minus daily minimum temperature as response variable also included the temperature of the old growth forest.

The difference between daytime plantation and old growth forest temperature decreased linearly with plantation age, at a rate of −0.420 °C / year (95% CI −0.590 - −0.250, figure 1a). Conversely, the nighttime temperature difference increased over time at a rate of 0.133 °C / year (95% CI 0.0853 – 0.180, figure 1a). The difference in daytime humidity between the plantations and old growth forest increased with time. The rate of change decreased with time, which is reflected by the smooth term having an effective df of 1.75 (figure 1b). For 5 year old plantations, the difference in humidity increased with a rate of 4.6% / year (95% CI 1.95 – 3.03), which dropped to 2.88 % / year (95% CI 0.83 – 4.92 % / year) at plantation age 15. The difference in nighttime humidity did not change with plantation age, with the 95% CI of the slope equalling −0.262 – 0.170 % / year (figure 1b). The diurnal temperature range – the difference between daily maximum and minimum temperature – decreased linearly with plantation age, with a slope of −0.476 (95 % CI −0.693 - −0.259, figure 1c).

**Figure 1.**
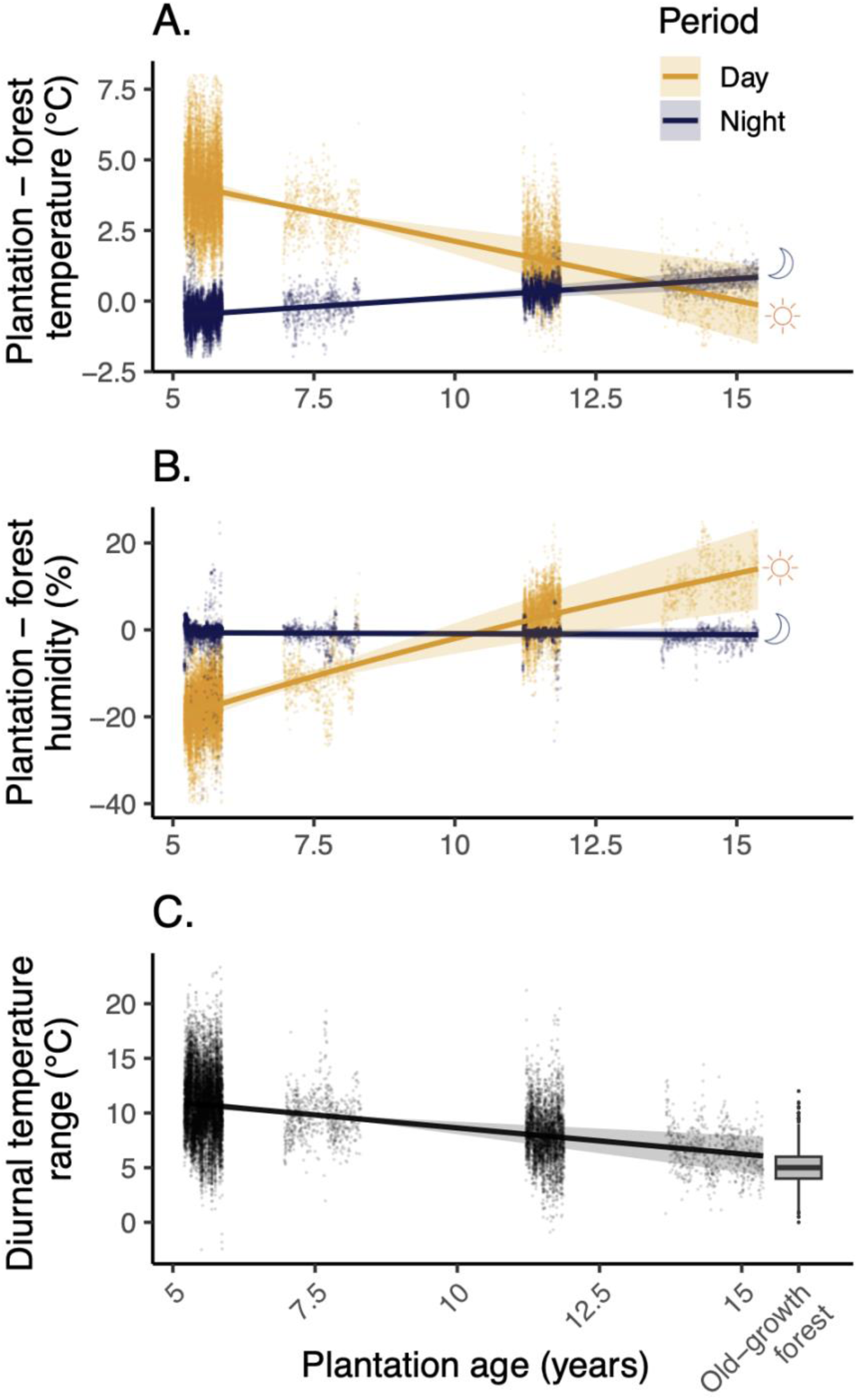
The difference in temperature (A) and relative humidity (B) between oil palm plantations and a nearby old-growth forest reserve, and the diurnal temperature range of the oil palm (C), as a function of plantation age, Differences are displayed separately for the daytime and nighttime. The curves show the smooths of Generalised Addiditive Mixed Models with their 95% confidence intervals. The scatter plots depict partial residuals. The temperature values were standardised for elevation and old-growth forest temperature using the model estimates. We used the elevation of the old-growth forest (432 m) and the mean temperature throughout the study (24.2°C for daytime data, 22.5°C for nighttime data). The boxplot in plot C gives the diurnal temperature range of the old-growth forest throughout the study period.

### Biotic data

The best models for ant and spider abundances included plantation age as a linear explanatory variable. Including distance to nearest forest did not improve the models (table S2). Ant abundance decreased with plantation age (slope = −0.11 ± SE 0.03, z = −3.5, p < 0.001, figure 2A and B), while spider abundance increased with plantation age (slope = 0.87 ± SE 0.27, z = 3.2, p = 0.001).

**Figure 2.**
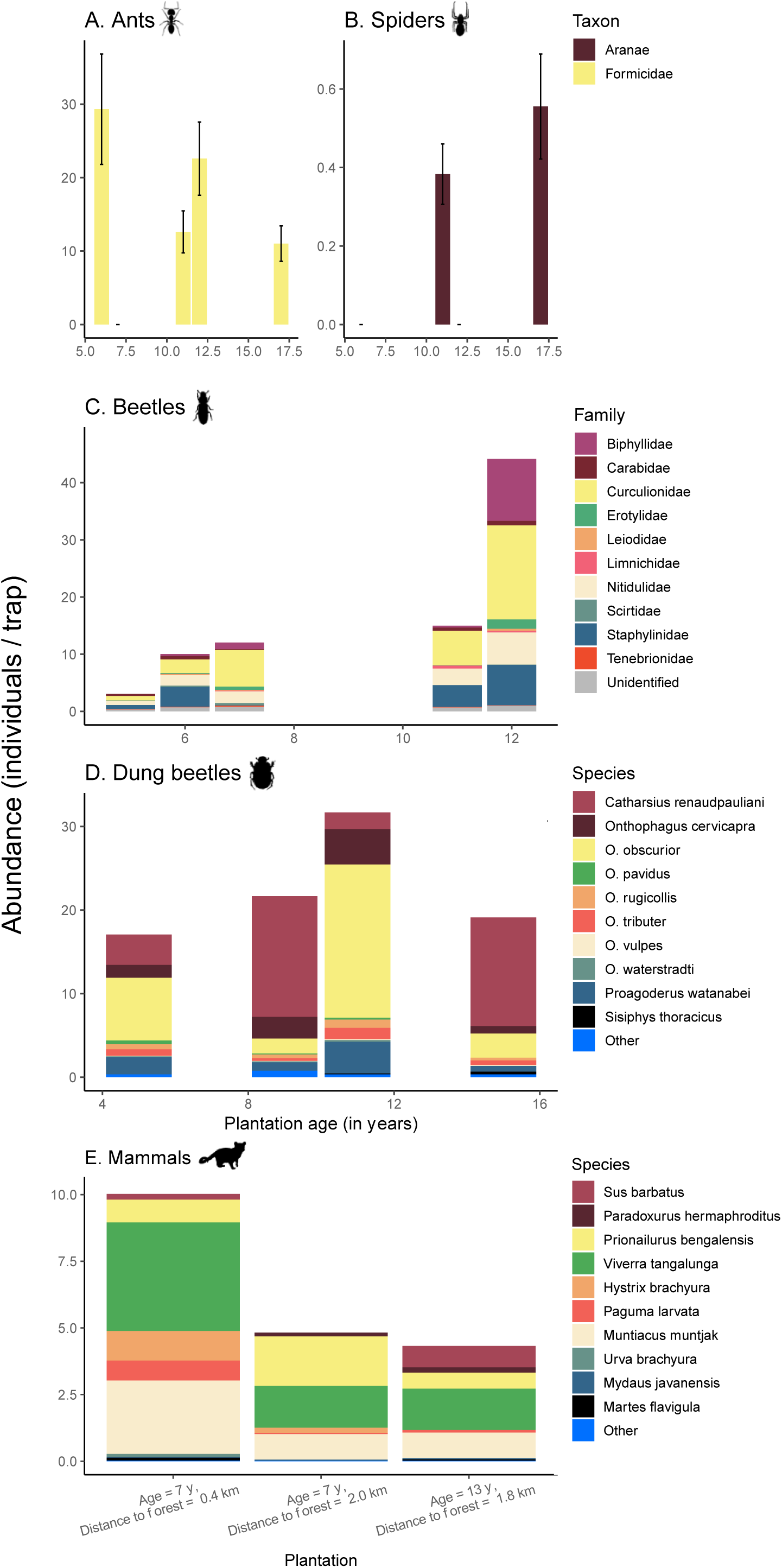
The abundance and community composition of different taxa over a chronosequence of oil palm plantation ages. For ants, spiders, beetle families, and dung beetle species (A - D), this is depicted for all sampled plantation ages. For mammals, it is displayed for all three plantations (E). Error bars denote standard errors.

Beetle abundance increased quadratically with plantation age (figure 2C, linear term = 10.1 ± SE 1.5, z = 6.6, p < 0.001, quadratic term = −2.3 ± SE 1.0, z = −2.24, p = 0.025). The multivariate model indicated that community composition significantly changed with plantation age (Dev_192,1_ = 191.3, p < 0.001) and distance to forest (Dev_192,1_ = 105.4, p = 0.002). Post-hoc univariate tests revealed that this was caused by the Biphyllidae (Dev_192,1_ = 38.4, p=0.001), Curculionidae (Dev_192,1_ = 39.6, p= 0.001), Erotylidae (Dev_192,1_ = 11.4, p = 0.036), Limnichidae (Dev_192,1_ = 14.4, p = 0.006), Nitidulidae (Dev_192,1_ = 23.9, p = 0.001), Staphylinidae (Dev_192,1_ = 17.8, p = 0.005) families increasing in abundance with plantation age, and the Staphylinidae family decreasing with distance to forest (Dev_192,1_ = 15.8, p = 0.041). Beetle family richness increased between plantation ages 5 and 7, but stabilised after this (figure 3).

**Figure 3.**
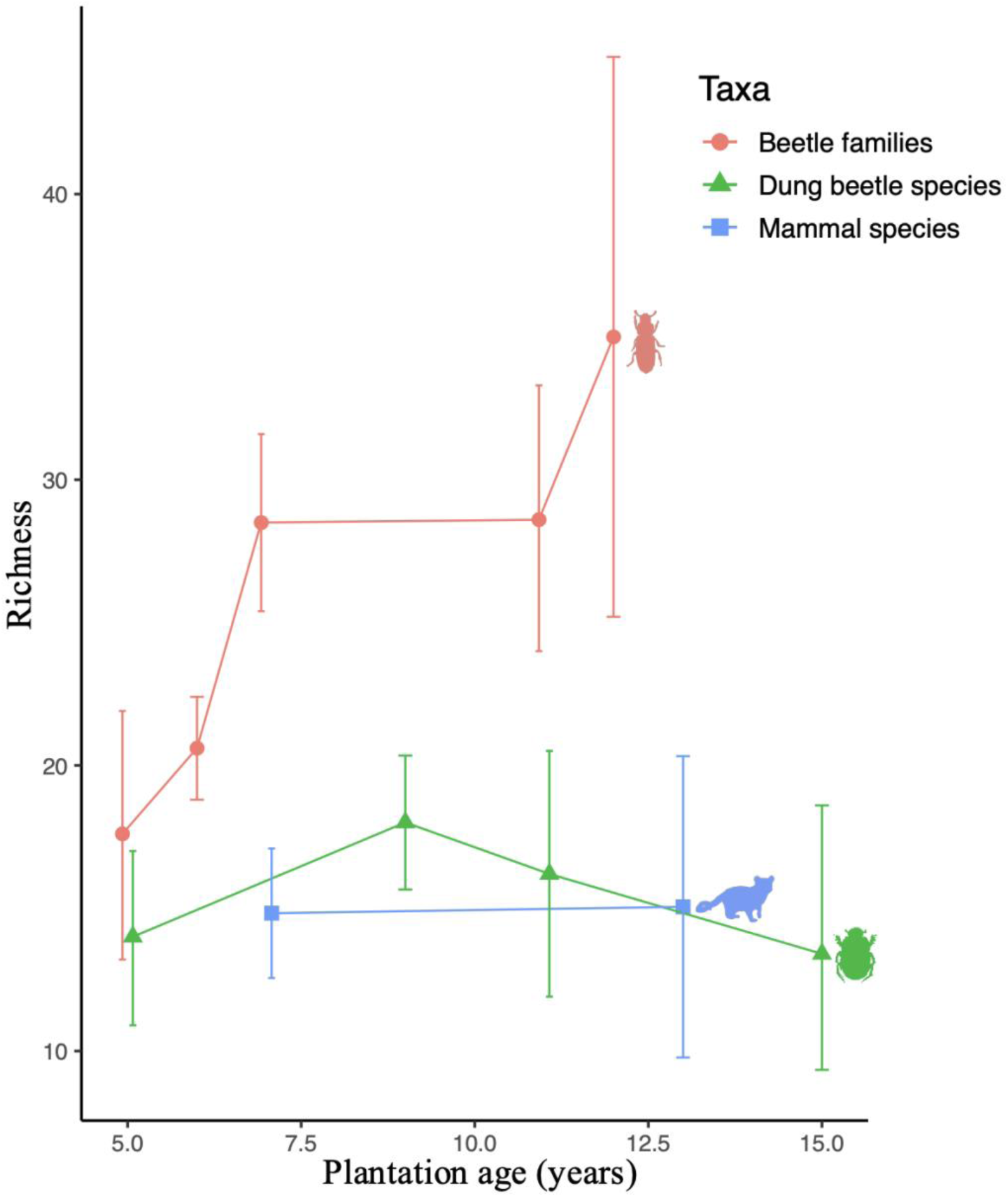
The richness of mammal and dung-beetle species, and beetle families, as a function of oil palm plantation age. Error bars denote 84% confidence intervals.

The model that best fit dung beetle abundance only included plantation age (table S2), although it was not a significant predictor (slope = 0.031 ± 0.023, z = 1.31, p = 0.19, figure 2D). Community composition, however, was significantly affected by both plantation age (Dev_52,1_ = 31.70, p = 0.014) and distance to forest (Dev_51,1_ = 68.3, p = 0.001). Post-hoc univariate tests indicated that this significant result was not caused by significant changes in the abundance of individual species. Dung beetle species richness did not change with plantation age, as revealed by the species accumulation curves of different plantation ages (figure 3).

The best fitting model for mammal abundance did not include plantation age (table S2), while distance to forest was a significant predictor, being negatively linked to mammal abundance (slope = −0.52 ± 0.10, z = −5.1, p < 0.001). Community composition was significantly affected by both distance to forest (Dev_66,1_ = 32.78, p = 0.003) and plantation age (Dev_67,1_ = 96.82, p = 0.001, figure 8A). The significance of the distance to forest variable was caused by red muntjac *Muntiacus muntjak* (Dev_66,1_= 12.914, p = 0.02), Malayan civet *Viverra tangalunga* (Dev_66,1_= 16.197, p = 0.01), masked palm civet *Paguma larvata* (Dev_66,1_= 9.461 p = 0.02), Malayan porcupine *Hystrix brachyura* (Dev_66,1_= 14.422, p = 0.02), short tailed mongoose *Urva brachyura* (Dev_66,1_= 8.193, p = 0.02), pig-tailed macaque *Macaca nemestrina* (Dev_66,1_= 10.115, p = 0.017), and spiny rat *Maxomys* sp. (Dev_66,1_= 7.786, p = 0.02), all declining with increasing distance to forest. No single species was significantly affected by plantation age. The mammals’ species accumulation curves showed that species richness did not differ between the plantations of different ages (figure 3).

## Discussion

Our results show that the microclimate of oil palm plantations becomes less harsh over time, and that this could make them more hospitable to some taxa. The microclimate is increasingly buffered, as the temperature drops, daily temperature fluctuations decrease, and humidity increases with plantation age. As such, the microclimate of the plantation increasingly resembled that of the nearby old-growth forest over time. Animal’s responses to plantation age varied between taxa. Spider and beetle abundances increased, ant abundance decreased, and mammal and dung-beetle abundances stayed constant, while the diversity of beetles in general increased. The degree to which animals are limited by plantations’ microclimate therefore appears to differ between taxa.

The more buffered microclimate of older plantations is likely due to their greater canopy cover and height (Jucker et al., 2018; Hardwick et al., 2015). This leads to a decrease in day-time temperatures as plantations age, since more sunlight is blocked by the canopy. Conversely, nighttime temperatures increase, since more of the day’s heat is trapped under the canopy, whereas the heat quickly escapes from the more bare, young plantations. In secondary forests, the microclimate similarly becomes more buffered over time, as the habitat recovers and canopy cover increases (González del Pliego et al., 2016). Hardwick et al., (2015) found that in addition to canopy cover, higher canopies also increase temperature buffering, because higher canopies allow less vertical mixing between ground and canopy, resulting in lower ground temperatures. Correspondingly, Luskin & Potts (2011) showed that 22-year-old oil palm plantations were colder and more humid than 8-year-old plantations, and that nighttime temperatures between forests and plantations did not differ. Plantations at the ages 5 to 7 experienced lower nighttime temperatures than forests, but the temperature difference between forests and plantations had disappeared at year 8. In the daytime, the temperature difference between forests and plantations continued to decrease between the ages 5 and 15 – indicating that the microclimate of oil palm plantations changes even after a decade into their lifecycle.

These continued changes in the microclimate of plantations partly explain the positive relationship between plantation age and spider and beetle abundance, and beetle diversity. The high temperatures of young plantations may make them inhospitable for many invertebrates (García-Robledo et al., 2016), thus limiting their abundance and diversity. As plantations age, and their microclimate increasingly resembles forests’, fewer species will be constrained by their thermal limits. These findings are largely congruent with the only other study that looked at invertebrate abundance relative to oil palm age, although that study focussed on second generation oil palms (Pashkevich et al., 2021): understory invertebrate abundance was higher in first generation plantations aged between 31 and 33 years, compared to 1, 3, and 8-year-old second generation oil palm plantations.

Besides the altered microclimate, the rise in yield could drive the observed increase of the abundance of several beetle families. Oil palm yield peaks around 10 years after planting, while biomass continually increases (Ling, 2012). Consequently, palms produce more inflorescences around this age, which could in turn boost pollinator abundance. We observed an increase in the abundance of the weevil family (Curculionidae), which contains oil palm’s most important pollinator, *Elaeidobius kamerunicus (Kouakou et al., 2018)*. As fruit production increases, the abundance of rotting fruit also increases, as more kernels spill. This could drive the increase of sap beetles (Nitidulidae), since many Nitidulidae species, including *Carpophilus mutilatus*, the most common Nitidulid in oil palm plantations, feed on rotting fruit (Nor Atikah et al., 2019). Older, bigger palms are also likely to more frequently produce new and bigger fronds. Fronds usually pruned, stacked, and left to decay in the plantation. This does not only provide an additional microhabitat, it also provides a food source. The decaying organic matter could in turn explain the rise in the beetle families Biphyllidae and Erotylidae. These families contain a high number of saprophagous species (Węgrzynowicz, 2015; Leschen & Buckley, 2007; Beutel & Leschen, 2016), which feed on this matter directly, and fungivores, which feed on the fungi that grow on decaying matter. This rise in fruit production, frond production, and decaying organic matter, therefore provides a clear mechanism by which plantation age, and improved yields could positively affect the biodiversity of plantations. Razak et al. (2020) demonstrated a positive link between the yield and bird-diversity of palm oil plantations, which could correlate with the increase in the abundance and diversity of beetles that we observed.

Our results show that ant abundance decreased with plantation age. This could be caused by initial high abundances of disturbance- and heat-tolerant ants, which become less abundant as the plantations recover. Plantations cool over time, which could cause the abundance of these heat tolerant ants to drop, to a degree that the increase in less heat tolerant, more specialist species, could not compensate for. Pashkevich et al. (2021) similarly found higher ant abundances in newly replanted oil palm plantations, compared to 3 year-old, 8-year-old, and 31 to 33-year-old plantations. Higher ant abundances have also been observed in disturbed habitats versus undisturbed habitats in the United States (Graham et al., 2004) and Malaysia (Luke et al., 2014).

Mammal abundance and diversity were not affected by plantation age. This is likely because as endotherms, mammals are not as restricted by the higher temperatures in young plantations (Clarke, 2014). Mammals might instead be limited by other aspects of the plantations, such as shelter and resource availability (Azhar et al., 2014). Needs such as these would explain why distance to forest was a more important predictor of mammal abundance than plantation age, and might indicate that mammals only venture short distances into plantations, while still depending on nearby forest habitat (Yue et al., 2015; Wearn et al., 2019).

The stability of mammal diversity over time could explain why dung beetle diversity and abundance also stayed constant. Dung beetles depend on mammals for food and breeding, and a strong link between mammal and dung beetle abundance and diversity has been well documented (Fuzessy et al., 2021; Culot et al., 2013; Raine et al., 2018). Despite the more favourable microclimate of older plantations, low mammal abundances are likely limiting the abundance of dung beetles.

We have shown that oil palm plantations undergo significant biotic and abiotic changes, even far into their lifecycle. Beetle abundance, for example, still increased between the 12^th^ and 16^th^ year since planting. Plantation age should therefore be taken into account when considering oil palm’s impact on biodiversity and climate (e.g. McAlpine et al., 2018; Sabajo et al., 2017). Correspondingly, snapshot biodiversity surveys of oil palm plantations cannot give a complete view of the biodiversity of plantations. This might also partially explain some of the inconsistency between the results of previous studies on the plantations’ biodiversity (Savilaakso et al., 2014). For example, Hashim, Jusoh & Nasir (2010) found no difference in ant abundance between mangrove forests and oil palm plantations, the youngest of which were 7-years old, while ant abundance was drastically lower in 14- to 18-year-old plantations compared to rainforests in peninsular Malaysia (Fayle et al., 2010). It could also help explain why smallholder plantations generally support higher biodiversity than large-scale plantations (Azhar et al., 2017). Smallholders tend to consist of palms of different stand-ages, whereas large scale industrial plantations do not have this diversity of stand-ages (Azhar et al., 2013).

As a result, large scale plantations are less structurally diverse than small-holders, which could result in a lower biodiversity (Azhar et al., 2015).

Our findings imply that habitat heterogeneity in oil palm dominated landscapes could be significantly improved by incorporating different stand-ages. Since the community composition of all taxa studied in this research changed significantly with plantation age, allowing landscape level variance in stand-age would increase overall biodiversity. As such, when replanting oil palms, it would benefit biodiversity to maintain strips of older palms (Luskin & Potts, 2011). This would ensure that some suitable habitat, for example for the beetle families that only occurred in older plantations, would be more likely to subsist. Such strips could potentially also be used to connect patches of forest habitat.

We have demonstrated that oil palm plantations are by no means static over time – their microclimate changes continually, even deep into their life cycle, and some taxa follow this trend. While our study focused solely on oil palm, this is likely the case for more types of tree plantation, such as rubber and timber plantations, since they follow a similar trajectory. However, few studies have attempted to make chronosequences of the microclimate and biodiversity of agricultural areas. We therefore still lack a comprehensive understanding of how their biodiversity and microclimate develops as crops age, including some widespread tree plantations such as rubber and eucalyptus. Future research should therefore focus on making chronosequences for other types of plantations, as well as for oil palm beyond the ages looked at in the current study. This can help us better understand the impact of agriculture on biodiversity, and guide landscape planning.

## Data availability statement

We intend to make all data publicly available on Zenodo.

## Author contributions

*Conceptualisation*: TAdL, RME, *Data curation*: all authors, *Formal analysis:* TAdL, *Methodology*: All authors, *Supervision:* RME, *Writing—review and editing*: All authors

## Acknowledgements

We thank the Sime Darby Foundation, the Sabah Foundation, Benta Wawasan, and the Sabah Forestry Department for granting permission to conduct fieldwork. We thank the South East Asian Rainforest Research Partnership for logistically supporting this project, and all research assistants involved with data collection. Research was funded by the Sime Darby Foundation. TAdL was supported by Imperial College London’s President’s PhD scholarship. EMS acknowledges funding from a British Ecological Society Small Ecological Project Grant, No.: 3256/4035, the Varley-Gradwell Travelling Fellowship in Insect Ecology, and the UK Natural Environment Research Council grant (NE/K016407/1). Dung beetle samples were collected under Permit No. EPU Ruj. UPE: 40/200/19/2712 and SaBC access licence number JKM/MBS.1000-2/2(381) and project number RS 302 under the Royal Society SEARRP. We thank A. Chung, R. Nilus, S.A. Mann, and J.M. Sawang for the support in collecting and coordinating the collection of the data.

