## Supplementary material for "Stand age drives changes in the biodiversity and microclimate of oil palm plantations": S1, S2, Table S1, S2

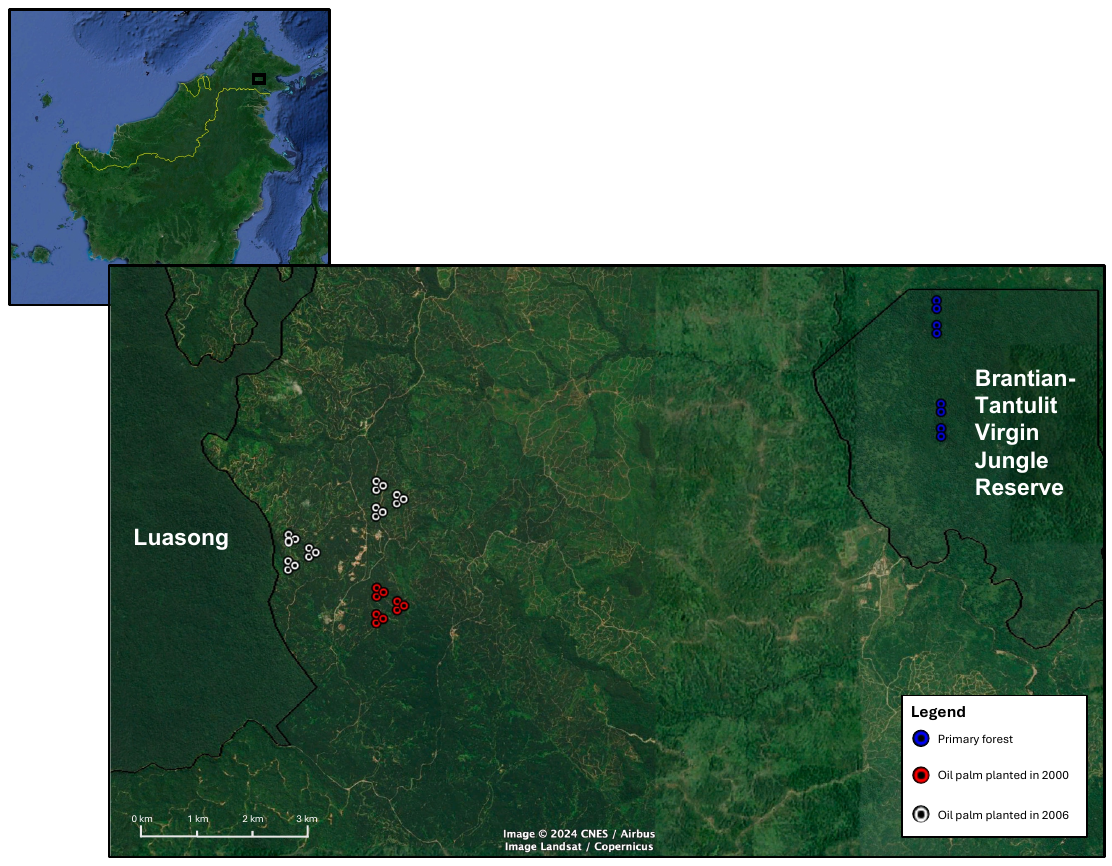

**Figure S1. A map of the sampling sites used in this study, and its location on the island of Borneo. Black lines indicate the borders between forests and oil palm plantations. Map adapted from Google Earth. Dots represent the second order sampling points – meaning that each dot forms the centre of a smaller triangle of first-order sampling points.**

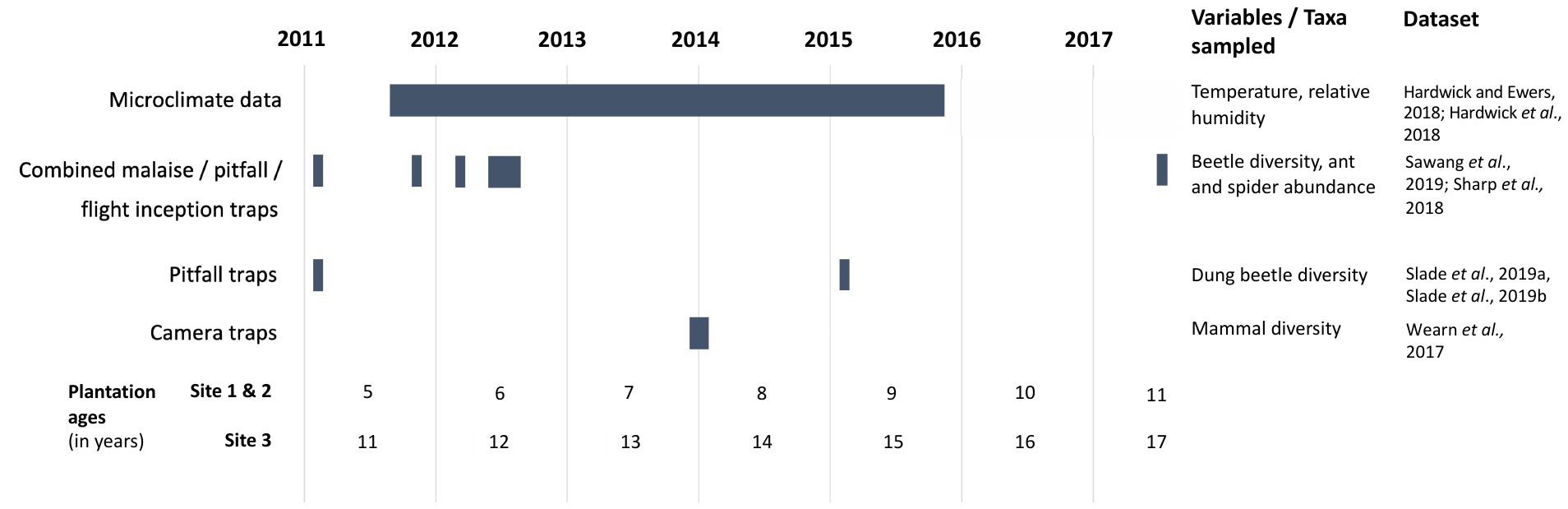

**Figure S2. A gantt chart showing the timing of the sampling campaigns used in this study, when they were conducted, which variables were measured, and the corresponding plantation ages.**

**Table S1. The results underpinning the selection of Generalised Additive Mixed Effects Models, describing how plantations microclimate changed as a function of plantation age.**

| **Mode** | **Included explanatory variables** | **Degrees of freedom** | **AIC** |
| --- | --- | --- | --- |
| Daytime temperature | Plantation age + Elevation + Old growth forest temperature | 12 | 45924.26 |
|  | Plantation age + Elevation | 11 | 47076.96 |
|  | Plantation age + Elevation + s(Old growth forest temperature) |  |  |
| Nighttime temperature | Plantation age + Elevation + Old growth forest temperature | 12 | 12077.82 |
|  | Plantation age + Elevation | 11 | 12581.63 |
|  | Plantation age + Elevation + s(Old growth forest temperature) | 13 | 11648.92 |
| Daytime relative humidity | Plantation age + Elevation + Old growth forest humidity | 12 | 91489.74 |
|  | Plantation age + Elevation | 11 | 92159.78 |
|  | Plantation age + Elevation + Old growth forest humidity + Old growth forest temperature | 13 | 90946.22 |
|  | Plantation age + Elevation + s(Old growth forest humidity) + s(Old growth forest temperature) | 15 | 89516.66 |
| Nighttime relative humidity | Plantation age + Elevation + Old growth forest humidity | 13 | 64738.11 |
|  | Plantation age + Elevation | 11 | 65483.87 |
|  | Plantation age + Elevation + Old growth forest humidity + Old growth forest temperature | 14 | 64702.18 |
|  | Plantation age + Elevation + s(Old growth forest humidity) + s(Old growth forest temperature) | 16 | 64689.70 |
| Diurnal temperature range | Plantation age + old growth forest temperature | 11 | 71219.37 |
|  | Plantation age | 10 | 72494.45 |
|  | Plantation age + s(Old growth forest humidity) | 12 | 70971.70 |

**Table S2. The model selection results of the negative binomial generalised mixed effects models, setting out the abundance of different taxa against several predictor variables.**

| **Taxon** | **Included explanatory variables** | **Degrees of freedom** | **AIC** |
| --- | --- | --- | --- |
| Mammals | Plantation age + distance to forest | 5 | 388.1236 |
|  | Plantation age | 4 | 396.7297 |
|  | Distance to forest | 4 | 387.0304 |
|  | Plantation age • Distance to forest | 6 | 390.0791 |
|  | Distance to forest^2^ | 5 | 384.4085 |
| Beetles | Plantation age + distance to forest | 8 | 2071.300 |
|  | Plantation age | 7 | 2069.356 |
|  | Distance to forest | 7 | 2147.209 |
|  | Plantation age • Distance to forest | 9 | 2073.282 |
|  | Distance to forest^2^ | 8 | 2143.986 |
|  | Plantation age^2^ | 8 | 2071.128 |
| Dung beetles | Plantation age + distance to forest | 5 | 418.1451 |
|  | Plantation age | 4 | 416.9705 |
|  | Distance to forest | 4 | 417.2670 |
|  | Plantation age • Distance to forest | 6 | 417.7475 |
|  | Distance to forest^2^ | 5 | 414.6766 |
|  | Plantation age^2^ | 5 | 418.2909 |
| Ants | Plantation age + distance to forest | 9 | 1659.285 |
|  | Plantation age | 8 | 1658.211 |
|  | Distance to forest | 8 | 1666.789 |
|  | Plantation age • Distance to forest | 10 | 1656.398 |
|  | Distance to forest^2^ | 9 | 1658.888 |
|  | Plantation age^2^ | 9 | 1668.405 |
| Spiders | Plantation age + distance to forest | 9 | 291.7881 |
|  | Plantation age | 8 | 289.9326 |
|  | Distance to forest | 8 | 336.6151 |
|  | Plantation age • Distance to forest | 10 | 293.7719 |
|  | Distance to forest^2^ | 9 | 335.3526 |
|  | Plantation age^2^ | 9 | 291.6869 |
